# Positive coping supports children’s emotional wellness: Behavioral evidence and neuroendocrine mechanisms

**DOI:** 10.1101/2023.02.19.526965

**Authors:** Ting Tian, Boxuan Chen, Yuyao Zhao, Hongyao Gao, Menglu Chen, Ying He, Jiahua Xu, Min Jiang, Bingsen Xiong, Shaozheng Qin

## Abstract

A positive coping style is recognized as a stable disposition to foster emotional wellness and resilience, enabling an adaptive process of assessing and dealing with environmental challenges. Such an adaptive process is believed to rely on a nuanced interplay of the hippocampal system and the primary stress hormone cortisol activity. As a hallmark of diurnal cortisol rhythm, cortisol awakening response (CAR) is sensitive to upcoming stress and subserves the preparation of the hippocampal system for rapid behavioral adaption. Yet, little is known about how the hippocampal system and CAR contribute to the merit of positive coping on emotional wellness. By two studies, we investigate the effects of positive coping on children’s emotional wellness and CAR, as well as longitudinal changes in hippocampal-neocortical functional systems involved in emotional processing. Behaviorally, positive coping predicted better emotional regulation ability, but lower anxiety and lower response caution in emotional decision-making. At the endocrine and neurocognitive level, positive coping was associated with greater CAR, which further predicted higher connectivity of the hippocampus with ventrolateral prefrontal cortex (vlPFC) and stimulus-sensitive neocortex one year later. Furthermore, CAR mediated an indirect association between positive coping and longitudinal increases in hippocampal-neocortical connectivity. Positive coping and CAR together could account for the maturity of vlPFC through longitudinal changes in hippocampal-neocortical connectivity. Overall, our findings suggest a cognitive-neuroendocrinal framework in which positive coping shapes hippocampal-neocortical maturation via stress hormone response to support emotional wellness.

**Significance:** The role of the hippocampal system in regulating stress response is well recognized, but its contribution to emotional well-being is not yet understood. Here we show that the protective effects of positive coping on emotional well-being are contingent on two factors: the cortisol awakening response (CAR), which is sensitive to upcoming stress, and hippocampal development. We found that positive coping practices promoted emotional wellness, enhanced emotional decision-making and increased CAR in young children. Longitudinal neuroimaging analysis revealed that positive coping-related CAR predicted greater hippocampal connectivity with stimulus-sensitive neocortex one year later. Importantly, CAR acted as a mediator of the promotive influence of positive coping on the longitudinal development of hippocampal-neocortical connectivity, which contributed to the maturity of prefrontal control systems. Our findings emphasize the importance of hippocampal-neocortical development in resilient coping and emotional wellness.

## Introduction

The accelerating pace of lifestyle in our contemporary society puts forward high requirements on our ability to rapidly learn and flexibly cope with ever-changing environmental demands, which poses a challenge to one’s mental health. A positive coping style is defined as the positive thoughts and strategies that an individual uses to manage internal and external demands in response to stress (1). Such coping strategies and behaviors can help individuals bounce back quickly from stressful events (2). Over the past decades, resilient coping is increasingly recognized as a stable disposition to buffer against stress susceptibility and allows an adaptive process of assessing stress and dealing with challenges to foster emotional wellness (3). School-age childhood is recognized as a sensitive period not only for early onset of various mental illnesses, but also for rapid development of the human brain, cognitive and affective functions. The importance of positive coping for maintaining emotional well-being has been well recognized at a behavioral level, however, the neurobiological mechanisms underlying positive coping from a developmental perspective are still in their infancy.

Positive coping has profound impacts on human psychosocial functioning and adaptive stress responses accompanied by the activity of stress-sensitive neuromodulatory systems (4). The stress bucket model posits that a positive coping style is like the tap of a bucket and it can buffer the pressure flexibly, otherwise stress response would overflow and then develop into mental disorders like anxiety (5). A more positive coping style, as a long-formed strategy of dealing with stressors, is closely related to plenty of mental cognitive resources (6) and is the integration of how individuals adapt to the ever-changing environment. Indeed, positive coping has a close link with emotional wellness and executive function (7). For example, individuals who tend to positively cope with things are able to recognize emotions better (8) and therefore benefit emotion regulation (9). Computational modeling of moment-to-moment decisive responses to negative emotional stimuli is regarded as a useful approach for uncovering latent cognitive processes involved in emotional decision-making, including drift rate and decision threshold measures that reflect the quality of evidence accumulation and response caution when making decisions(REFs). Few studies to date, however, have addressed the underlying latent cognitive processes of how positive coping affects emotional wellness in childhood. By another aspect, research in psychology and endocrinology has provided robust evidence that the influence of positive coping on stress responses and mental health outcomes involves an interplay of the behavioral, cognitive and endocrinal systems (10, 11). In particular, the Hypothalamus-Pituitary-Adrenal axis (HPA axis) produces “stress hormone” cortisol to control the response to stress (12, 13) and regulates neural activity subtly to maintain homeostasis. However, even few studies (14) have directly explored the relationship between coping style and cortisol response. Therefore, it remains a gap between assumption and empirical research on the effects of positive coping on individual emotional wellness from a view of latent computational mechanisms as well as the endocrinal processes.

Recent neurobiological models have suggested that stress-sensitive cortisol, acting as one of the mediators, actively shapes neuronal activity and communication of brain networks critical for the perception and regulation of emotional stimuli and events (15, 16). The stress generation model proposes that each individual’s cognitive and behavioral characteristics, such as belief and negative coping style, are the vulnerable factors of psychopathology (17, 18) especially for affective disorders (19). Such abnormalities have been linked to complex alterations in the HPA-axis activity (20). Cortisol awakening response (CAR) is a hallmark of HPA-axis functioning(21), which is a burst of cortisol within 30-45 minutes after morning awakening and could prepare and store energy for upcoming stress according to the brain preparedness theory of CAR (22). Among the limbic brain system, the hippocampus is key to regulating stress responses, assessing and adapting to environmental changes, and dealing with daily challenges (23, 24). Importantly, the hippocampus plays an essential role in fine-tuning cortisol levels (25, 26) due to its dense collection of mineralocorticoid receptors (MRs) and sparse distribution of glucocorticoid receptors (GRs) (27, 28), and these two receptors could bind with cortisol to maintain homeostasis balance and meet the environmental demands. A sub-optimal level of CAR may lead to structural damage in the hippocampus and hippocampal functional connectivity (29). Recent studies show that early life stress could affect children’s limbic-prefrontal connections through cortisol secretion (30, 31), suggesting the potential mediation effect of endocrinal cortisol response on psychosocial functioning and hippocampal development. However, whether and how positive coping along with CAR shapes hippocampal functioning in development remains open.

The neuroaffective association with resilient coping has been extensively approached from a perspective of functional localization in the amygdala across species (32–34), while ignoring the role of the hippocampus in the processing of emotional information, whose engagement is critical for emotional learning, memory (35, 36) and resilient regulation of stress response (37). For example, a series of experiments in one recent study reveal that the hippocampus adaptively updates negative memories by finding positive meaning in negative events (38), highlighting the significance of hippocampal engagement in positive coping regulating emotional memory. Critically, the hippocampus develops rapidly during childhood (39) and its development is especially sensitive to stress (40) and the alteration of stress hormone cortisol (41), thus brain maturation of the hippocampus is significant to maintain stable emotional states. Besides, the hippocampus has an inhibition effect on the HPA-axis system and such a negative feedback function is also modulated by the regulation of prefrontal activity (24). The functional coordination between the hippocampus and ventrolateral prefrontal cortex (vlPFC), as a critical neural pathway for top-down regulation of emotional memory (42) and involved in modulation of cortisol levels (24), shows an increasing trend with age (43), indicating the plasticity of hippocampus-vlPFC functional development. From a perspective of child brain development, insufficient attention has been paid to the psychophysiological interaction effects of positive coping and CAR on the longitudinal adaptation of the hippocampus during emotion processing. Although positive coping, cortisol response and hippocampal systems have been separately investigated by many isolated studies, it still remains unknown about how positive coping works in concert with CAR to shape the development of the hippocampal-neocortical systems in emotion processing during childhood.

To address the above open questions, we conducted two studies to investigate the behavioral, endocrine and neurocognitive correlates of how positive coping promotes children’s emotional wellness. In Study 1, we examined the beneficial effects of positive coping on emotional regulation skills, trait and state anxiety as well as CAR in 89 school-age children. In Study 2, we further investigated the effects of positive coping and CAR on hippocampal-neocortical functional maturation involved in emotional processing by using longitudinal fMRI spanning over two years in 34 school-age children. The CAR was obtained by the difference in cortisol values between the awakening point and 30 minutes after awakening at Time-1. Computational modeling was implemented to uncover latent processes underlying decision-making during emotion processing. Univariate, multivariate and connectivity analyses were conducted to examine the development of hippocampal functional coupling. Mediation analyses were applied to test the mediation effect of CAR on the relationship between positive coping and the development of hippocampal connectivity. Based on the above theoretical models of resilient coping and stress-sensitive cortisol activity in humans, we hypothesized that positive coping would promote children’s emotional regulation skills and emotional wellness such as reduced anxiety, along with higher CAR. Moreover, positive coping would impact the hippocampus-neocortical maturation through CAR in childhood.

## Methods and Materials

### Participants

A total of 89 school-aged children participated in this longitudinal study (Time-1 age: 6-12, 8.23 ± 1.47 years old, 36 girls and 53 boys). A subsample of 34 children was invited back one year later (Time-2 age: 9.06 ± 1.50 years old; duration: 12.60 ± 2.02 months, **Table S1** in **Supplement**). None of them reported a history of neurological/psychiatric disorders and medical treatment in the past 6 months. A written informed consent form was obtained from the children’s parents/legal guardians and their willingness to participate from the children. All protocols were approved by the local institutional review board under the standards of the Declaration of Helsinki.

### Positive coping style

Positive coping style was measured by one dimension of the Simplified Coping Style Questionnaire (SCSQ) (44). The SCSQ, based on the “Ways of Coping” questionnaire, assesses two dimensions of coping strategies including active and passive coping strategies with self-reported 20 items. The active coping dimension mainly reflects active coping strategies individuals use when encountering stress, such as “trying to see things in as good a way as possible” and “identifying several different ways to solve problems.” Responses are given on a four-point Likert scale (0 = never; 3 = very often). The SCSQ scores reflect participants’ coping style preferences, with a higher score indicating a higher possibility that one would adopt the relevant coping style (45). The instrument of SCSQ has been widely used, with high reliability and validity (Cronbach’s α ranging between 0.90 and 0.92) (44).

### Emotion regulation and anxiety assessment

Emotion regulation was assessed by a 10-item self-reported Emotion Regulation Questionnaire (ERQ), which is widely used in measuring emotion regulation ability (46). ERQ involves two dimensions of emotion regulation ability, corresponding to two different emotion regulation strategies, i.e., cognitive reappraisal (6 items) and expressive suppression (4 items). Children were asked to rate their feelings and expressions of emotion on a 7-point Likert scale from strongly disagree to strongly agree. The ERQ showed good internal consistency and its test-retest reliability within 2 months is 0.7 (46). Only the cognitive reappraisal dimension of emotion regulation was significantly associated with positive coping in our data, therefore we used the reappraisal dimension as an indicator of children’s emotion regulation ability throughout the manuscript.

Anxiety was measured by using a self-reported State-Trait Anxiety Inventory (STAI) (47), with a well-trained assessor’s assistance. The STAI consists of 40 items, of which the first 20 items are tested for the state anxiety subscale and the last 20 items are tested for the trait anxiety subscale. Children were instructed to rate how frequently each item occurred in their lives on a 4-point Likert scale, with 1 - “not at all” and 4 - “very much”. Notably, children were asked to rate the most appropriate feelings for state anxiety at the moment, while the trait anxiety subscale was required to rate the most frequent feelings. The total score of state anxiety and trait anxiety ranged from 20 to 80, and this anxiety measurement is widely used to assess children’s anxiety by a previous study (48). Emotion regulation scores and anxiety scores over ± 3 standard deviations (SD) were considered outliers and excluded from further analyses.

### Salivary Cortisol Sampling and Analysis

Before cortisol sampling, children and their parents were given verbal and written instructions on how to collect cortisol to ensure effective accurate cortisol assessment. The salivary cortisol samples were collected by Salivette collection devices with their parent’s assistance (Sarstedt, Germany). All participants were asked not to brush their teeth, rinse their mouth, smoke, drink or eat for at least one hour before collection and not to smoke, drink or do strenuous exercise the night before. Five salivary cortisol samples were collected for each child at the following time points: pre-bedtime basal cortisol level before sleep (around 10-11 pm, N0), immediately (N1), 15 minutes (N2), 30 minutes (N3), and 60 minutes (N4) after awakening the next morning (**Table S2** in **Supplement**). For each cortisol sampling, children were required to record on the tube label the precise time at which they completed their salivary cortisol sampling. After collection, all the cortisol samples were brought back to the experimenter and stored at −80° C until assay. Salivary samples of children who reported any sickness, any hormonal medication or menstruation (only for girls) were excluded from further analyses.

Saliva samples were thawed and centrifuged at 3200 revolutions per minute (RPM) for 10 minutes. The concentration of cortisol was analyzed by electrochemiluminescence immunoassay with a sensitivity of 0.500 nmol/L and a standard range of 0.5-1750 nmol/L for determination (48). Each cortisol sample was naturally log-transformed to ensure a normal distribution. The rising cortisol level within 30 minutes after awakening (R30) was computed by the cortisol level at 30 minutes after awakening (N3) minus the cortisol level at awakening point (N1) (49), and the R30 values higher than ±3 SD were excluded from further analyses. R30, as one of the indicators of CAR, is thought to reflect an accelerating activity of the HPA-axis system during the transition from sleep to wakefulness.

### FMRI Task Procedures

In a six-minute emotion-matching task scanning session, children were instructed to view a trio of negative facial expressions or mosaic ovals in either emotion or control condition respectively. They were asked to identify which stimulus in the bottom row displayed the same emotional category or direction as the target stimulus in the top row (**Figure 2**). This task is considered a widely used negative emotion paradigm that consisted of 5 emotion and 5 control blocks with 6 trials each (50, 51). By contrasting the emotion versus control condition, this paradigm can assess emotion-related brain responses to emotional reactivity (52, 53).

### Hierarchical drift-diffusion model (DDM)

The DDM was widely implemented to model individual decision dynamics in the 2-choice decision-making tasks. This model posits that there is a dynamic and fluctuating process of information collection behind individual behavioral decisions, and when the collected information exceeds a certain threshold, decisions such as individual response time (RT) and accuracy (ACC) will be generated (54). The Hierarchical drift-diffusion model (HDDM) could model the individual-level decision parameters including the decision threshold (a), drift rate (v) and non-decision time (t), therefore the group statistical analyses are available. The decision threshold indicates the boundary of information collection or the decision criteria, and the individual tends to choose a certain option of 2-choice when the collected information reaches such a boundary. The drift rate determines the speed of deciding evidence accumulation. The non-decision time reflects other factors affecting decision time before the decision process, like stimulus encoding and executing responses.

HDDM were fitted to trial-by-trial RTs for responses under emotion condition and control condition emotional matching task for each child, and trials with RTs < 0.05 second were treated as outliers (55). We run multiple models consisting of different decision parameters, therefore resulting in 7 models including a, v, t, av, at, vt, avt. The model comparison was conducted with the deviance information criterion (DIC), whose lower value means better model fitting (56) (**Figure S1** in **Supplement**). The winning model in our study was avt model and the Markov Chain Monte Carlo (MCMC) method was used in this model to generate 20,000 samples from the joint posterior parameter distribution and discarded the first 2000 samples as burn-in (57). The model convergence was further evaluated by Gelman-Rubin statistic and the r-hat values of all parameters were close to 1.0 and less than 1.1 (54), indicating good convergence. The three parameters a, v, t from the winner model were extracted for subsequent analyses.

### Image Data Acquisition and Preprocessing

This study utilized Siemens 3T scanner (TIM Trio, Erlangen, Germany) to acquire whole-brain functional images, using a 12-channel head coil and a T2*-sensitive echo-planar imaging sequence (33 axial slices, 4-mm slice thickness, 0.6-mm gap, 2000-ms repetition time, 30-ms echo time, 90 flip angles, voxel size 3*3*4 mm^3^, field of view 200*200 mm^2^). Functional images were preprocessed through SPM12 (https://www.fil.ion.ucl.ac.uk/spm/software/spm12/). Considering MR signal stabilization, the first five volumes of the resting scan and the first four volumes of the emotional task scan were discarded. The remaining images were corrected for slice acquisition timing, realigned for head motion correction, spatially normalized into the Montreal Neurological Institute (MNI) space, resampled into 2-mm isotropic voxels, and smoothed by convolving a 6-mm isotropic three-dimensional Gaussian kernel.

### Univariate general linear model (GLM) analysis

To assess task-evoked neural activity during the emotion matching task, the emotion condition and control condition were modeled as two separate regressors and convolved with the canonical hemodynamic response function (HRF) implemented in SPM12. To remove the head motion-related artifacts, each participant’s Friston-24 head-motion parameters were included as nuisance covariates. The imaging data were high-pass filtered with a cutoff of 1/128 Hz and were corrected for serial correlations using a first-order autoregressive model (AR (1)) in the GLM framework. Contrast images were calculated at the individual level to identify brain regions with significant activation under the emotion condition versus control condition. Next, the individual-level contrast images were submitted to group-level multiple regression analyses with children’s R30 as the covariate of interest while child age and sex as nuisance covariates. Significant clusters were determined at a height threshold of *p* < .001 and an extent threshold of *p* < .05 corrected using 3dClustSim (58) for multiple corrections. Parameter estimates were then extracted from significant clusters to characterize activation patterns.

### Generalized psychophysiological interaction (PPI)

The generalized PPI method (59) was used to examine the task-evoked functional connectivity of the hippocampus, which was defined as a seed from the Automated Anatomical Labeling (AAL) template. The mean time series of all voxels within the hippocampus were computed and subsequently deconvolved to estimate the hippocampal neural activity. Next, two individual-level PPI regressors under the emotion condition regressor and control condition were calculated by multiplying the estimated hippocampal neural activity with a dummy vector indicating the task condition and therefore forming two psychophysiological interaction vectors. Then, the two PPI vectors were further convolved with a canonical HRF to form two PPI regressors of interest. The data were also high-pass filtered with a cutoff of 1/128 Hz and corrected for serial correlations using AR (1). In addition, each participant’s Friston-24 head-motion parameters and task-induced neural activity were included as nuisance covariates. Individual-level contrast images of PPI effects were submitted into second-level multiple regression analyses with R30 as a covariate of interest while child age and sex as nuisance covariates to reveal the significant hippocampal connectivity related to R30. Significant clusters were determined by a height threshold of *p* < .001 and an extent threshold of *p* < .05 with 3dClustSim multiple comparisons. Mean parameter estimates of significant clusters were then extracted to show their relation with R30.

### Mediation Analysis

Mediation models were constructed to examine whether CAR mediated the relation between positive coping and hippocampal connectivity. Besides, a chain mediation model was built to test whether positive coping affected the vlPFC maturation sequentially through R30 and the development of hippocampus-fusiform connectivity. The mediation models and statistical tests were conducted through PROCESS in SPSS version 21 (IBM Corp., Armonk, NY) (60). The magnitude and the significance of all models were calculated using the 5000 bias-corrected bootstrapping resampling approach (61), which produced a 95% confidence interval (CI) for the direct effect and indirect effect. If the 95% CI of the indirect effect did not include zero, the indirect effect was considered significant.

## Results

### Positive coping supports children’s emotional wellness and emotional decision-making dynamics

We first examined whether and how positive coping supports emotional wellness by focusing on emotional regulation ability, trait and state anxiety as well as emotional decision-making dynamics in Study 1. Pearson correlation analyses showed that positive coping was significantly correlated with higher reappraisal scores of emotion regulation (*r* = 0.38, *p* < 0.01, **Figure 1A**), and negatively with trait anxiety (*r* = −0.49, *p* < 0.01, **Figure 1B**) and state anxiety (*r* = −0.32, *p* < 0.01, **Figure S2A** in **Supplement**). Moreover, we investigated whether positive coping could predict emotional decision-making dynamics when performing a negative facial expression matching task (**Figure 1C**). We applied the HDDM (**Figure 1D**) to uncover the latent processes of emotional decision-making dynamics for each trial of emotion condition and control condition at both Time-1 and Time-2. This analysis revealed that higher positive coping was correlated with lower decision threshold (parameter a) under emotion condition at both Time 1 (**Figure S2B** in **Supplement**) and Time-2 (both *r* = −0.44, *p* < 0.05, **Figure 1E**). These results indicate the beneficial effects of a positive coping style on children’s emotional wellness, with increased emotional regulation ability, attenuated trait and state anxiety as well as lower decision threshold during negative emotion processing.

**Figure 1.**
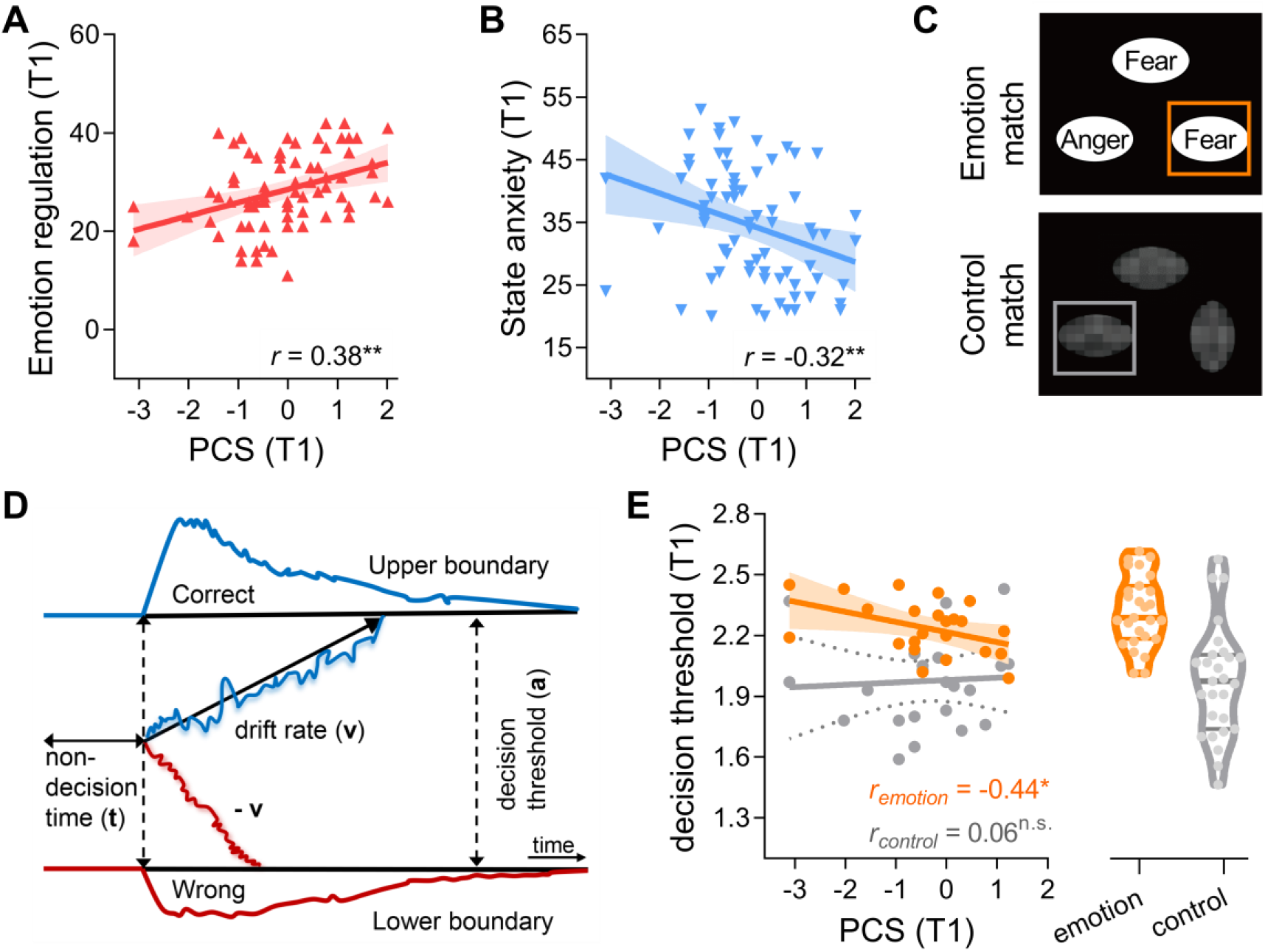
Positive coping promotes positive mental states and predicts behavioral decision dynamics. (**A**) Positive coping was positively related to emotion regulation ability in reappraisal dimension. (**B**) higher positive coping was associated with lower trait anxiety score. (**C**) The simplified graphical representation of the emotion matching paradigm, during which children were asked to match the same emotional category or shape direction with the target emotional face or shape. (**D**) Schematic illustration of HDDM with 3 model parameters: drift rate (v) indicates the speed of estimation during evidence accumulation until decision boundary or threshold (a) is reached. v is showed in blue lines for correct responses and red lines for wrong responses while a is showed by the vertical arrow on the right. The non-decision time (t) indicates the other factors affecting decision time, showing by the horizontal arrow on the left. (**E**) Positive coping predicts the decision boundary (a) under emotion condition at Time-2 but not for control condition. The right panel shows the data distribution of parameter a in two conditions. PCS, positive coping style; T1, Time-1; T2, Time-2; **, *p* < 0.01; *, *p* < 0.05; n.s., not significant.

### Positive coping increases CAR at the physiological hormonal level

Next, we examined the effects of positive coping on stress hormone response. We first evaluated the profile of morning cortisol levels at four-time points (i.e., N1, N2, N3 and N4). As shown in (**Figure 2A**), the cortisol level at N3 was significantly higher than N1 (*t* = 2.68, *p* < 0.01), indicating an effective CAR effect in healthy children. We then assessed CAR by computing an increase of cortisol = within post-awakening 30 minutes – that is, the difference between N1 and N3 time point, namely R30. Pearson correlation analysis revealed higher positive coping score predictive of greater R30 (*r* = 0.27, *p* = 0.02, N=72, **Figure 2B, Table 1**). Machine learning-based prediction analysis with a 10-fold cross-validation method revealed that greater positive coping predicted higher R30 (*r*_*(predicted, observed)*_ = 0.21, *p* = 0.02). These results indicate positive coping predictive of greater CAR in children, likely reflecting greater accelerating activity of the HPA-axis function in the morning.

**Figure 2.**
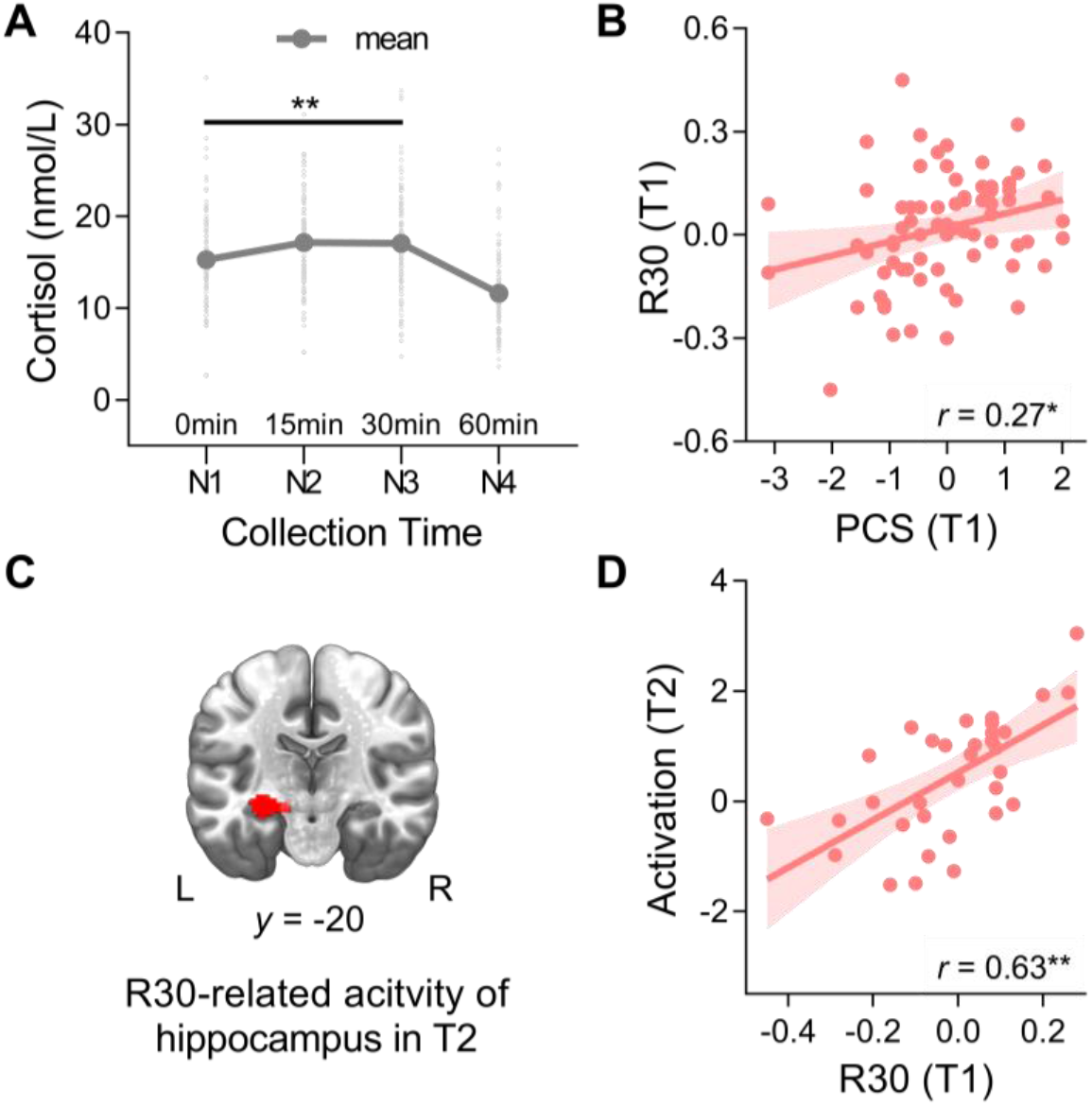
Positive coping increases CAR and predicts the hippocampal engagement at Time-2. (**A**) The dynamic curve of cortisol collected at four times: the awakening point in the morning (N1), 15 minutes (N2), 30 minutes (N3) and 60 minutes (N4) after awakening. Each light gray point represents a cortisol value of a participant at a specific time point and the dark gray thick line shows the mean cortisol changes of all children. The N3 was significantly higher than N1, indicating an effective CAR curve. (**B**) A positive correlation between positive coping and R30, which was calculated by the difference between N3 and N1 (i.e., N3-N1). (**C-D**) R30 was positively related to the activation in the hippocampus at Time-2. PCS, positive coping style; R30, response within 30 minutes; L, left; R, right; T1, Time-1; T2, Time-2; **, *p* < 0.01; *, *p*<0.05.

### Positive coping and CAR promote longitudinal changes in hippocampal functioning

More importantly for the question at issue, we examined whether and how positive coping and CAR shape brain systems involved in emotional processing. We conducted separate multiple regression analyses for task-invoked brain activity maps with positive coping and CAR as covariates of interest respectively when regressing out age and sex. The CAR-related regression analyses revealed significant clusters in the hippocampus (**Figure 2C-2D**) and other regions including the dorsolateral prefrontal cortex (dlPFC, **Figure S3, Table S3** in **Supplement**). Machine learning-based prediction analyses further confirmed that the first-year CAR positively correlated with task-invoked activation in the hippocampus (*r*_*(predicted, observed)*_ = 0.53, *p* = 0.002), which is what we are most interested in. Although no hippocampal effect was found in the positive coping-related regression analyses, we found that positive coping could predict the involvement of brain regions like the medial prefrontal cortex (mPFC, *r* = 0.59, *p* < 0.01) at Time-2 (**Figure S4** in **Supplement**). Together, the first-year CAR could predict the hippocampal activity one year later in the emotion-processing task.

### Positive coping modulates hippocampal functional connectivity one year later via CAR

In addition to hippocampal activity, hippocampal connectivity could provide information about the functional coupling of the hippocampus and other brain sites. To assess how positive coping-related CAR influences the hippocampal functional coupling at Time-2, we applied multiple regression analyses for the bilateral hippocampus (**Figure 3A**) seed-based functional connectivity maps at Time-2 with R30 (Time-1) as a covariate of interest while regressing out child age and sex. We found that CAR was significantly correlated to the hippocampus connectivity with fusiform (*r* = 0.58, *p* = 0.001, **Figure 3B**) and the ventrolateral prefrontal cortex (vlPFC, *r* = 0.45, *p* = 0.016, **Figure 3C, Figure S5, Table S4** in **Supplement**). A 10-fold cross-validation method also was used to examine whether R30 could predict the hippocampal connectivity at Time-2, and these analyses showed that higher R30 predicted increased hippocampus connectivity with fusiform (*r*_*(predicted, observed)*_ = 0.54, *p* = 0.003) as well as hippocampus connectivity with vlPFC (*r*_*(predicted, observed)*_ = 0.40, *p* = 0.01), suggesting robust predictive power of CAR on the hippocampal connectivity at Time-2.

**Figure 3.**
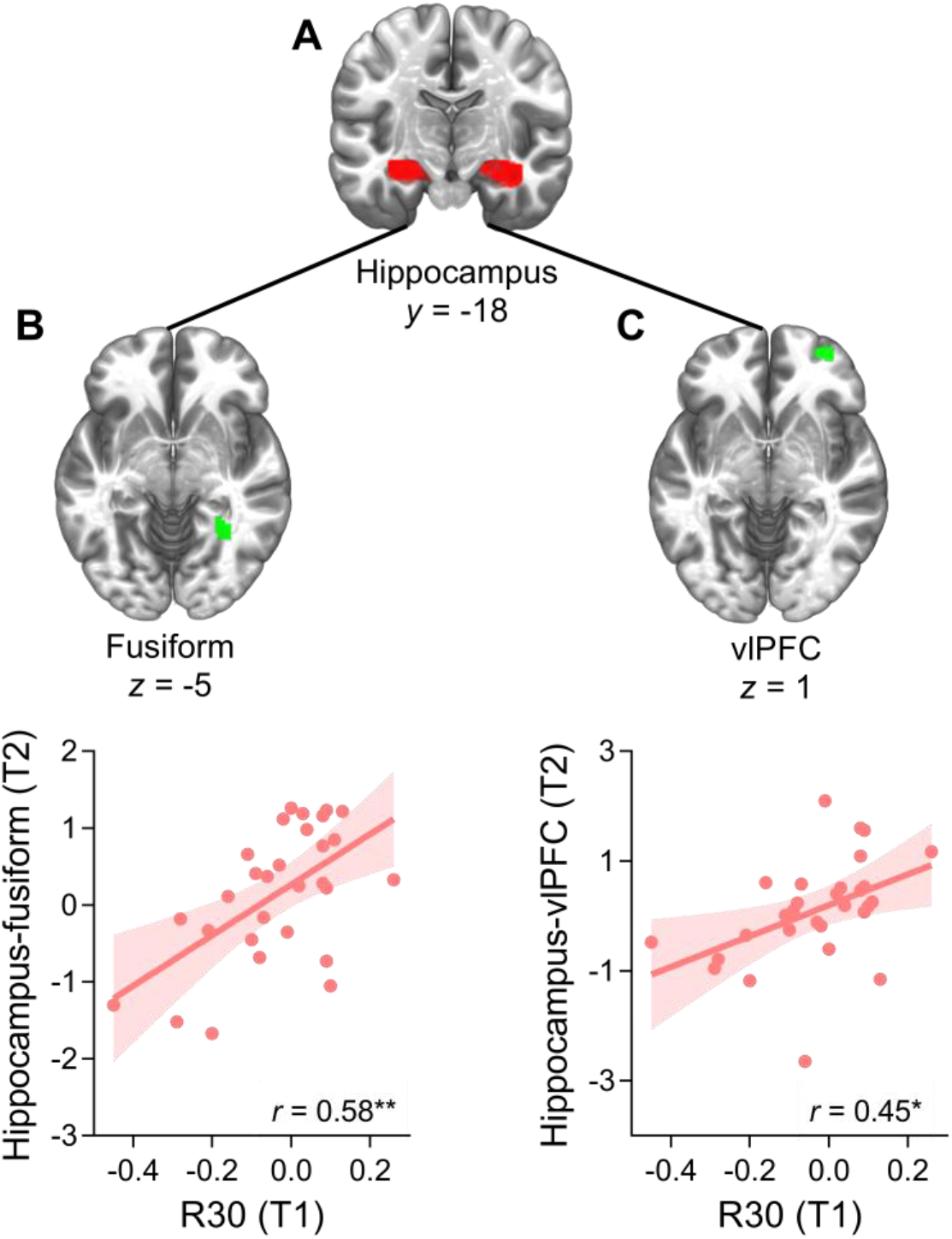
CAR predicts the hippocampal connectivity at Time-2. (**A**) A representative coronal slice of the seed region of bilateral hippocampus. (**B**) A positively significant relationship between the first-year R30 and the hippocampus-fusiform connectivity at Time-2, as well as (**C**) the relationship between R30 (Time-1) and the hippocampus-vlPFC connectivity at Time-2. R30, response within 30 minutes; T1, time 1; T2, time 2; vlPFC, ventrolateral prefrontal cortex; *, *p* < 0.05; **, *p* < 0.01.

To test whether positive coping affects hippocampal connectivity through CAR, we conducted mediation analyses to examine the mediation effects of CAR. These analyses revealed that positive coping predicted higher CAR at Time-1, which in turn accounts for hippocampal connectivity with the vlPFC (beta = 0.15, SE = 0.12, 95% CI = [0.023, 0.503]) as well as hippocampal connectivity with the fusiform gyrus (beta = 0.15, SE =0.11, 95% CI = [0.005, 0.426], **Figure S6** in **Supplement**). These results indicate that the CAR mediates an indirect relationship of positive coping with hippocampus-vlPFC at Time-2 and hippocampus-fusiform connectivity at Time-2.

### Positive coping-related CAR shapes hippocampal-neocortical functional maturation

We also investigated whether the CAR mediates the relationship between positive coping and the direct developmental changes in hippocampal connectivity over one-year interval, i.e., the change of hippocampal connectivity from Time-1 to Time-2. Mediation analyses were used to examine whether positive coping modulates the direct changes in hippocampal functional coupling through CAR. Mediation analyses revealed that CAR played a mediatory role in the indirect association between the first-year positive coping scores and changes in hippocampal-fusiform connectivity (i.e., Time2-Time1, beta = 0.16, SE = 0.10, 95% CI = [0.019, 0.404], **Figure 4A-4B, Figure S7-S8, Table S5** in **Supplement**), but not for hippocampus-vlPFC connectivity (**Figure S8-S9** in **Supplement**). Furthermore, we examined whether such an effect could predict longitudinal changes in the vlPFC (i.e., Time-2 minus Time-1) multivariate maturation index. We here defined the maturation index as the similarity of vlPFC multivoxel activity in children as relative to the mature adult brain, and the development of the vlPFC maturation index as the change in the vlPFC activity similarity between children and adults using Time-2 minus Time-1 (illustrated in **Figure 5A**). The correlation analysis revealed that R30 at Time-1 was positively related to longitudinal changes in vlPFC maturation index from Time-1 to Time-2 (**Figure 5B**). Both longitudinal changes in hippocampal-fusiform connectivity and vlPFC maturation index were z-transformed for further analysis. We then applied a chain mediation analysis and found that positive coping increased children’s R30 at Time-1, which subsequently accounted for greater longitudinal changes in hippocampal-fusiform connectivity (i.e., Time-2 minus Time-1) and finally promoted the change of vlPFC multivariate maturation index (beta = 0.04, SE = 0.04, 95% CI = [0.001, 0.170], **Figure 5C**). Finally, we used our hippocampus seed-based functional connectivity map to conduct decoder analysis which decodes and calculates the terms in Neurosynth that have the highest correlation coefficients with our hippocampal connectivity map. The image decoder allows to quantitatively and interactively compare the hippocampus functional connectivity map to the images database in Neurosynth (62). The decoder analyses showed that the hippocampal connectivity is positively related to the terms including working memory, tasks, memory, demands and load, while negatively related to the terms including emotional, theory mind, facial expressions, negative and mental states (**Figure 5D**), indicating the interplay of positive coping and CAR on meeting the environmental demands and reducing negative emotions. Taken together, these results reveal that positive coping may cause higher CAR in the endocrinal level to account for the development of lower-level hippocampal-fusiform connectivity and the development of higher-level vlPFC maturation.

**Figure 4.**
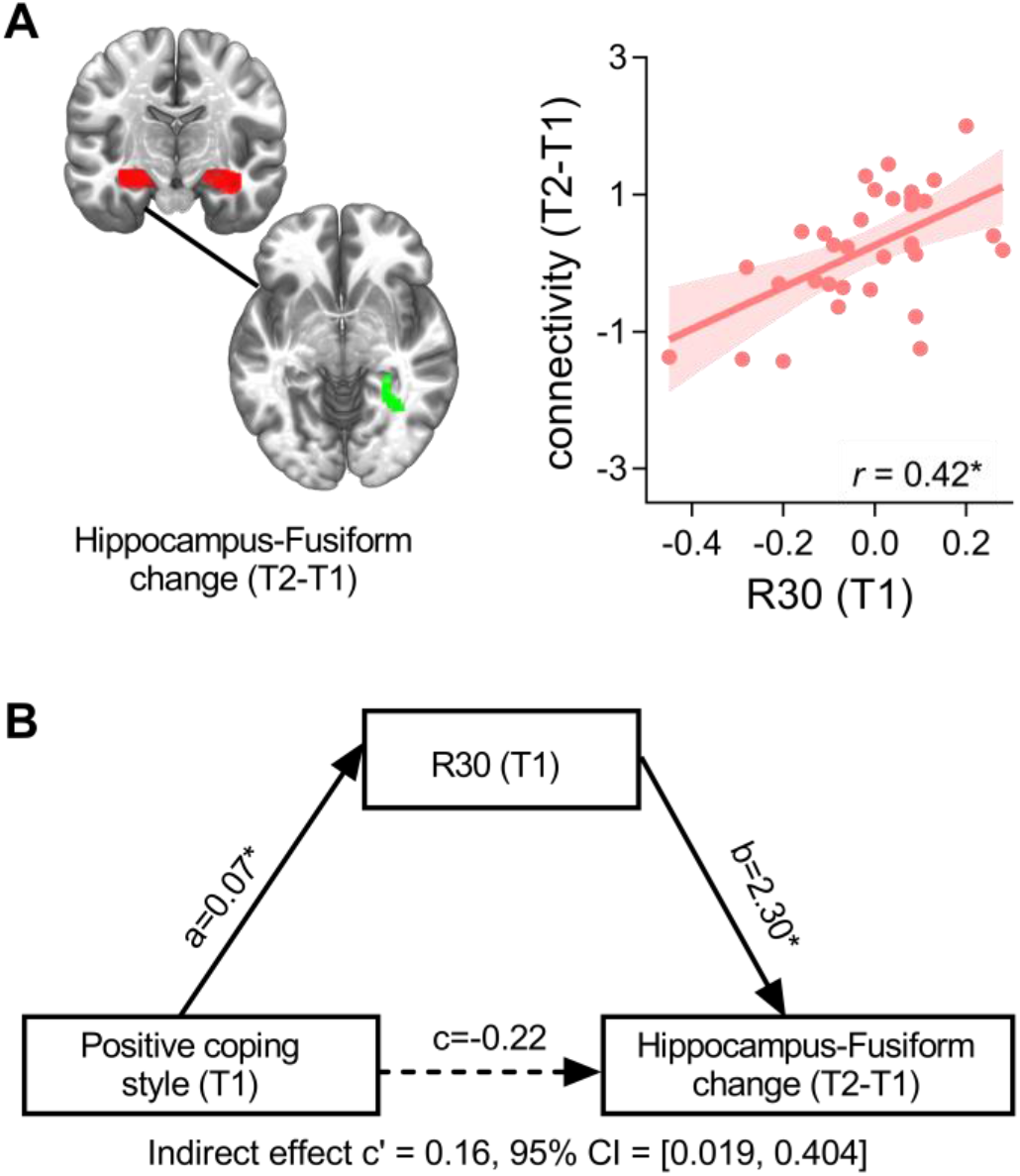
CAR mediates the relation between positive coping and the development of hippocampus-fusiform connectivity (Time2-Time1). (**A**) Left panel: R30 at Time-1 was significantly related to the change of the hippocampus-fusiform functional connectivity (Time2-Time1) with the bilateral hippocampus as the seed region (upper red brain area). Right panel: a scatter plot showing a positive relation between R30 and the change of hippocampus-fusiform connectivity over one year (Time2-Time1). (**B**) A mediation model indicating the mediation effect of R30 on the relation between positive coping and the connectivity change of the hippocampus-fusiform over two time points (Time2-Time1). T1, Time-1; T2, Time-2; R30, response within 30 minutes; *, *p* < 0.05.

**Figure 5.**
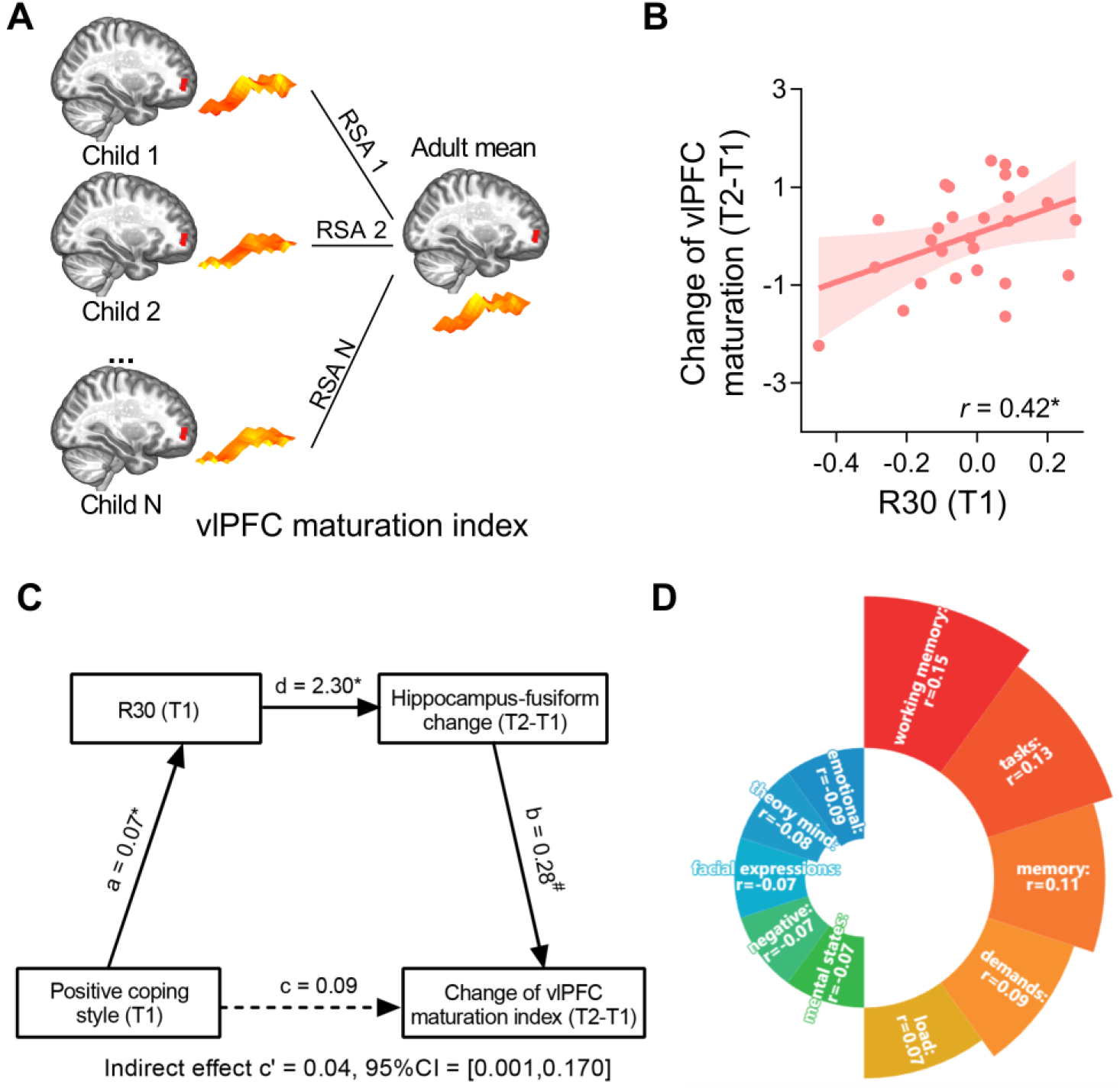
CAR mediates the relation between positive coping and the development of hippocampus-fusiform connectivity (Time2-Time1), which in turn predicts the vlPFC maturity change (Time2-Time1). (**A**) the diagram of vlPFC maturation index, which is calculated by the RSA in the vlPFC between children and adults. The change of vlPFC maturation index is the difference of the second-year RSA minus the first-year RSA between children and adults. (**B**) A significant association between R30 and the change of vlPFC maturation index (Time2-Time1). (**C**) A chain mediation model showing that positive coping affected R30 first, then affected the change of vlPFC maturation index over one year (Time2-Time1) through the change of hippocampus-fusiform functional connectivity (Time2-Time1). (**D**) The image decoder analyses using the bilateral hippocampus seed-based brain connectivity map. The five red areas mean the top positive terms that were positively correlated with the hippocampus connectivity map while the five blue areas mean the top terms that were negatively related to the hippocampus connectivity map. SA, representational similarity analysis; vlPFC, ventrolateral prefrontal cortex; R30, response within 30 minutes; T1, time 1; T2, time 2; *, *p* < 0.05; ^#^, *p* = 0.07.

## Discussion

By two studies, we investigated the behavioral, endocrine and neurocognitive substrates underlying how positive coping supports children’s emotional wellness. As expected, positive coping predicted better emotion regulation skills, lower anxiety and greater CAR. This paralleled greater hippocampal activity and increased hippocampal connectivity with stimulus-sensitive fusiform and vlPFC one year later. Critically, stress-sensitive CAR mediated an indirect relationship between positive coping and longitudinal changes in hippocampal-neocortical functional connectivity. Furthermore, positive coping-related CAR modulated the maturity of the vlPFC through a longitudinal increase in hippocampal-fusiform connectivity (Time-2 vs Time-1). Our findings suggest that greater CAR acts as a key mediator to scaffold the promotion effects of positive coping on hippocampal-neocortical functional coupling in support of emotional development in childhood.

### Positive coping supports emotional wellness, with better emotional regulation ability, reduced anxiety and lower response caution

Our results of positive coping improving children’s emotion regulation ability and decreasing anxiety indicate the promoting effects of positive coping on emotional competency and emotional wellness. Positive coping integrates wider processes involving cross-talk among regulation of stress response, cognitive control, and working memory (63, 64), therefore it may be closely linked with the adaptive process and key to flexibly responding to challenges. Firstly, the positive relationship between positive coping and emotion regulation possibly lies in the point that positive coping has broader dimensions than emotion regulation and shares certain similar features with emotion regulation (65, 66), including generating moderate responses to stressful circumstances and making efforts to regulate emotions (67). Children using a more positive coping strategy tend to look on the bright side of things and utilize a wealth of emotional skills, which accounts for a higher ability for emotional reappraisal. Secondly, our results showed that positive coping reduced both state anxiety and trait anxiety, which may be possibly related to the sub-components of positive coping revealed in previous studies, namely seeking social support (68) and adopting flexible ways to solve problems (69), and such results are consistent with the close link between positive coping and decreased risk for a variety of mental outcomes.

Interestingly, outcomes from our computational modeling of moment-to-moment decisive responses showed that positive coping predicted a lower decision threshold (i.e., model parameter a) one year later during a negative emotional decision-making task. This measure is believed to reflect the boundaries of information collection for evidence accumulation during which individuals more readily make a decisive choice when the collected information reaches such criteria. As pointed out in the previous review, positive coping is not only the heart of dealing with diverse challenges but also the core framework for executive control, especially in the psychosocial decision-making process (70). The positive coping strategy benefits the flexible allocation of attention resources and social information processing (71), therefore individuals with more positive coping may reflect to have more confidence to recognize emotional cues and tend to make corresponding decisive choices with less caution, which may account for the reliably profound effects of positive coping on decreasing the emotional decision boundaries. In a word, positive coping is a core component of flexible adaptation process and it benefits to improve children’s emotional wellbing and optimize the latent emotional decision-making process.

### Positive coping-related CAR predicts longitudinal hippocampal activity

In conjunction with behavioral effects, we found that higher positive coping correlated with greater CAR, which in turn predicted the longitudinal engagement of the hippocampus. CAR is an accelerating HPA-axis activity within post-awakening 30 minutes when switching from the sleep state at night to the wake state in the morning (72), and such a rapid rise in cortisol secretion is essential for the body to provide sufficient energy supply for metabolic consumptions and the environmental challenges (22, 73). Although lacking substantial direct evidence about the positive coping-CAR relation, our results are consistent with the positive relationship between resilience and CAR (14). Furthermore, results from our longitudinal neuroimaging revealed that more positive coping-related greater CAR could shape future hippocampal engagement. According to the brain preparedness theory of CAR, CAR could provide energy for upcoming events and prepare the brain for environmental demands, therefore greater CAR may mobilize sufficient resources for the hippocampal response to emotional events and stress response. Besides, the hippocampus gives negative feedback to the HPA-axis system and adjusts the glucocorticoid (cortisol in humans and corticosterone in rodents) to a moderate level (74, 75). Glucocorticoid modulates memory and learning through their binding with two types of corticosteroid receptors, including higher-affinity Mineralocorticoid receptors (MRs) which are greatly abundant in the hippocampus, and lower affinity Glucocorticoid receptors (GRs) which are widely distributed among diverse brain sites (27). This may mean that the predominantly fast occupation of MRs by cortiosol occurs in the hippocampus first and therefore it may impact the hippocampal regulation of emotional memory and memory updating. Our results that higher CAR predicts greater hippocampus activity one year later during the emotion processing task may reveal that the HPA-axis activity activates the hippocampus to retrieve emotional memories and reflect the hippocampal role in regulating the emotional response to negative stimuli (76). Moreover, this point may be supported by abundant evidence showing that excessive cortisol elevations would impair hippocampus-dependent memory performance and hippocampal anatomy, suggesting a critical link between CAR and hippocampal engagement (77, 78). In a word, positive coping-related CAR predicts longitudinal hippocampal activity, suggesting the hippocampal role in evaluating stressors and coping with future challenges under the HPA-axis regulation.

### Positive coping and CAR shape hippocampal-neocortical functional maturation facilitating executive control

Beyond regional activation in the hippocampus, we found that positive coping led to higher supportive CAR and modulated higher hippocampus connectivity with fusiform of lower-level sensory-motor area and vlPFC of higher-level prefrontal area under emotion condition versus control condition at the connectivity level, and such a connectivity pattern benefited individual executive control including working memory and responses to task demands (shown in Figure 5D). The functional coupling between the hippocampus and the fusiform is thought to be involved in socioemotional processing including processing facial expression (79), and these results that positive coping modulates the development of hippocampus-fusiform connectivity via CAR may mean that the higher positive coping and CAR collectively activate more hippocampal engagement with the sensory-visual system to actively process the information input of emotional faces. Furthermore, we found that CAR modulated higher hippocampus connectivity with higher-level vlPFC under emotion condition versus control condition, which is presumably because that children with higher CAR may have sufficient energy to support the hippocampal inhibition of the HPA-axis responses to socio-emotional stimuli (80), accompanied by the top-down regulation of vlPFC to the hippocampus (42). Such a similar pattern of higher hippocampus-vlPFC connectivity can be seen in responses to negative stimuli (42) and the process of episodic memory retrieval (81).

Although positive coping did not directly predict the hippocampus-vlPFC connectivity change (T2-T1) through greater CAR, we interestingly found that the interplay of positive coping and CAR affected vlPFC maturation through the hippocampus-fusiform connectivity change. This may imply that children with more positive coping and higher CAR might retrieve past emotion-related information to utilize in current emotional events. In other words, one would actively mobilize existing cognitive resources to cope with the current emotional stressors when receiving negative emotional stimuli. Such a phenomenon is revealed in resilient people that actively utilize positive emotions to exert effective control over stressful situations (82), also seen in interventions using positive memory to offset negative memory in a very recent neuroimaging study (38), suggesting individuals’ subjective initiative in emotion regulation, which is the core opinion of positive psychology theory (83). To sum up, the interplay of positive coping and CAR affects the vlPFC maturation via the development of hippocampus-fusiform connectivity, showing positive coping and CAR support children’s ability of emotion regulation in response to negative stimuli.

**In conclusion**, our study shows that positive coping has far-reaching promotion effects on children, including improving their emotional wellness, optimizing the emotional decision-making process, increasing CAR of the HPA-axis activity and facilitating the maturation of the hippocampus-neocortical functional systems which subserves executive control. Importantly, positive coping shapes the development of hippocampal function through CAR. Our findings provide insights into the promotive role of hippocampal-neocortical maturation in support of neuroendocrinal modulation, positive coping and emotional wellness. Overall, these findings advance our understanding of the developmental psychopathology framework and neuroendocrinal mechanisms underlying resilient coping.

