## Supplement for "Positive coping supports children’s emotional wellness: Behavioral evidence and neuroendocrine mechanisms"

#### *Supplemental Information*

#### Supplementary Tables

**Table S1. Participant demographics of Study 1 and Study 2**

| Study | *N* | Boys/Girls | Age Range | Average Age ± SD |
|---|---|---|---|---|
| Study 1 | 89 | 53/36 | 6-12 years | 8.28 ± 1.47 years |
| Study 2 | 34 | 22/12 | 6-12 years | 9.06 ± 1.50 years |

*N*, number of participants; SD, standard deviation.

**Table S2. The cortisol levels of four saliva cortisol samples in the morning.**

| Study | N1 ± SEM | N2 ± SEM | N3 ± SEM | N4 ± SEM |
|---|---|---|---|---|
| Study 1 | 15.27 ± 0.55 | 17.13 ± 0.57 | 17.06 ± 0.61 | 11.61 ± 0.50 |
| Study 2 | 17.08 ± 0.73 | 18.29 ± 1.09 | 17.16 ± 1.02 | 11.98 ± 0.79 |

Notes: cortisol levels in the awakening point (N1), post-awakening 15 minutes (N2), 30 minutes (N3), and 60 minutes (N4) in the morning.

**Table S3. The whole brain activations in the second year (T2) that were significantly related to children's R30.**

| **Target Regions** | **L/R** | **BA** | ***T* value [a]** | **MNI (x y z)** |
|---|---|---|---|---|
| Inferior Occipital Gyrus | R167 | 18 | 3.95 [a] | 38 -92 -6 |
| Supramarginal Gyrus | R417 | 40 | 4.58 [a] | 38 -48 34 |
| Hippocampus | L79 | 28 | 4.23 [b] | -28 -22 -12 |

[a] Clusters were thresholded with a height threshold of $p < 0.001$ and an extent threshold of $p < 0.05$ corrected for multiple comparisons. [b] Given our prior hypothesis, significant clusters in the hippocampus were determined at a height threshold of $p < 0.001$ and an extent threshold of $p < 0.05$ corrected for multiple comparisons with an anatomically defined bilateral hippocampus mask from Automated Anatomical Labeling (AAL) template. CMA, centromedial amygdala; BLA, basolateral amygdala; BA, Brodmann's area; MNI, Montreal Neurological Institute; -, no proper data.

**Table S4. Target regions of the hippocampus seed-based connectivity in the second year (T2) that were significantly related to children's R30.**

| Seed | Target Regions | L/R | BA | *T* value [a] | MNI (x y z) |
|---|---|---|---|---|---|
| | Inferior Occipital Gyrus | R167 | 18 | 3.95 [a] | 38 -92 -6 |
| | Inferior Parietal Gyrus | L121 | 40 | 3.97 [a] | -44 -48 36 |
| Hippocampus | Supramarginal Gyrus | R417 | 40 | 4.58 [a] | 38 -48 34 |
| | Superior Frontal Gyrus | R52 | 10 | 3.65 [b] | 28 56 -6 |
| | Fusiform | R50 | 19 | 4.72 [c] | 32 -52 -4 |

[a] Clusters were thresholded with a height threshold of $p < 0.001$ and an extent threshold of $p < 0.05$ corrected for multiple comparisons. [b] Given our prior hypothesis, significant clusters in the superior prefrontal cortex (vlPFC) were determined at a height threshold of $p < 0.001$ and an extent threshold of $p < 0.05$ corrected for multiple comparisons with an anatomically defined vlPFC mask or [c] face fusiform area (FFA) mask from Automated Anatomical Labeling (AAL) template. BA, Brodmann's area; MNI, Montreal Neurological Institute.

**Table S5. Target brain regions with significant changes in functional connectivity with the hippocampus seed (T2-T1) that were related to children's R30.**

| **Seed** | **Target Regions** | **L/R** | **BA** | ***T* value** [a] | **MNI (x y z)** |
|---|---|---|---|---|---|
| Hippocampus | Fusiform | R57 | 19 | 4.13 [a] | 32 -52 -4 |

[a] Given our prior hypothesis, significant clusters in the superior prefrontal cortex (vlPFC) were determined at a height threshold of $p < 0.001$ and an extent threshold of $p < 0.05$ corrected for multiple comparisons with an anatomically defined face fusiform area (FFA) mask from Automated Anatomical Labeling (AAL) template. BA, Brodmann's area; MNI, Montreal Neurological Institute.

### Supplementary Figures

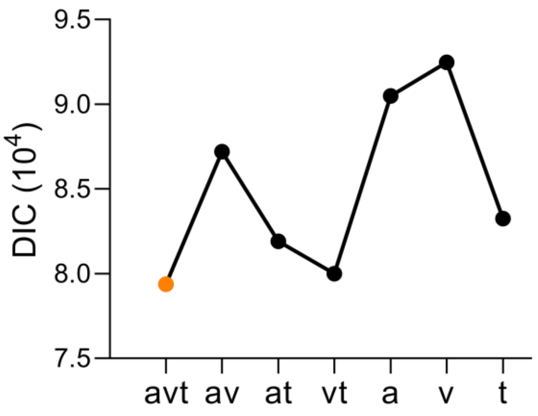


**Figure S1. The model comparisons of HDDM.** DIC differences between the best-fit model and the other model variants. The best model was avt model and consists of the boundary (a), drift rate (v), and non-decision time (t) during the 2-choice emotion matching task. DIC, deviance information criterion.

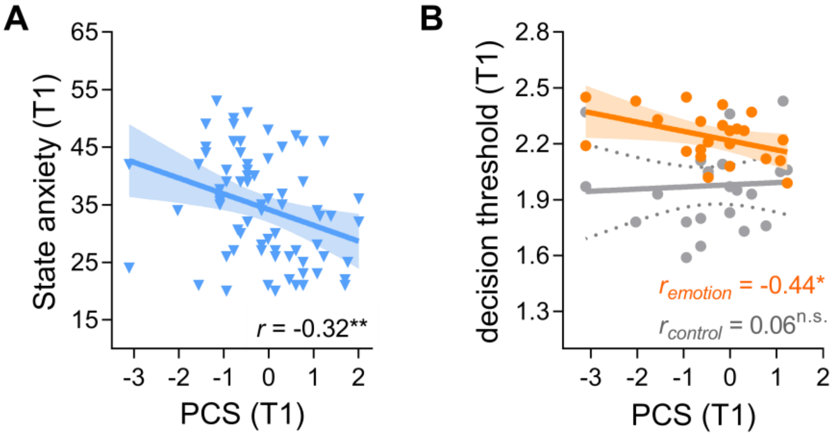


**Figure S2. Positive coping was negatively related to state anxiety at Time 1 and predicted decision threshold at Time-1.** PCS, positive coping style; **, $p <$ 0.01; *, $p < 0.05$; n.s., not significant.

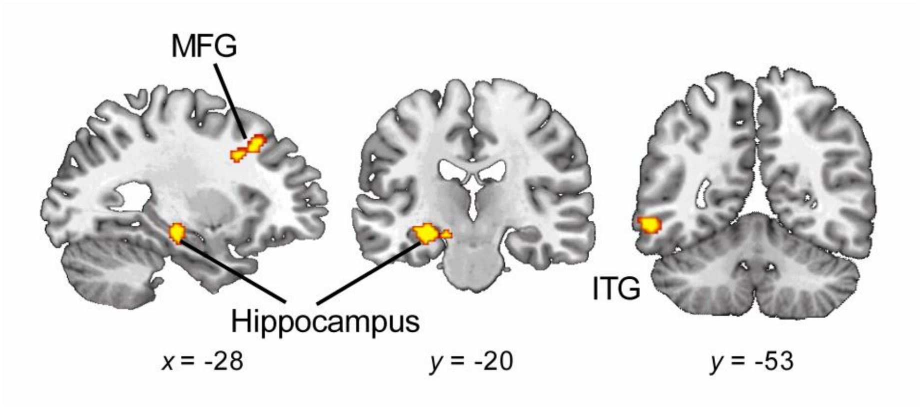


**Figure S3. The whole brain activations in the second year (T2) that were significantly related to children's R30.** MFG, middle prefrontal gyrus; ITG, inferior temporal gyrus; x means the sagittal view; y means the coronal view.

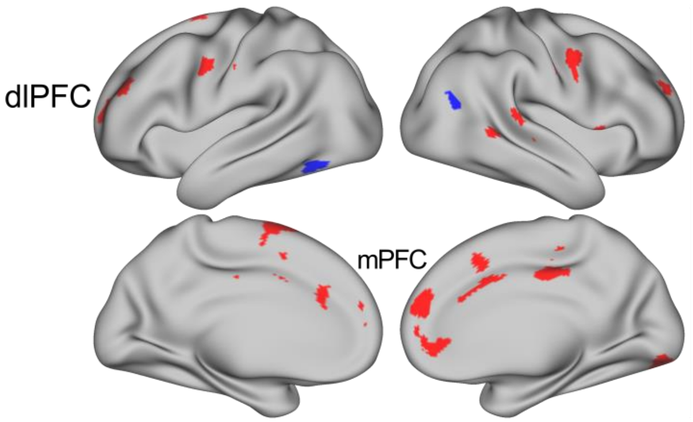


**Figure S4. The PCS-related whole brain clusters at Time-2.** dlPFC, dorsolateral prefrontal cortex; mPFC, medial prefrontal cortex.

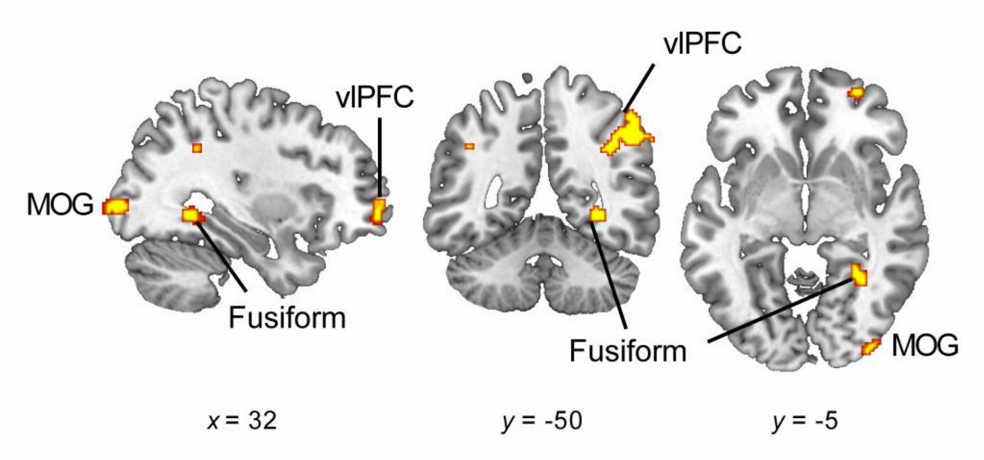


**Figure S5. The Target brain regions of the hippocampus seed-based connectivity in the second year (T2) that were significantly related to children's R30.** MOG, middle occipital gyrus; vlPFC, ventrolateral prefrontal cortex.

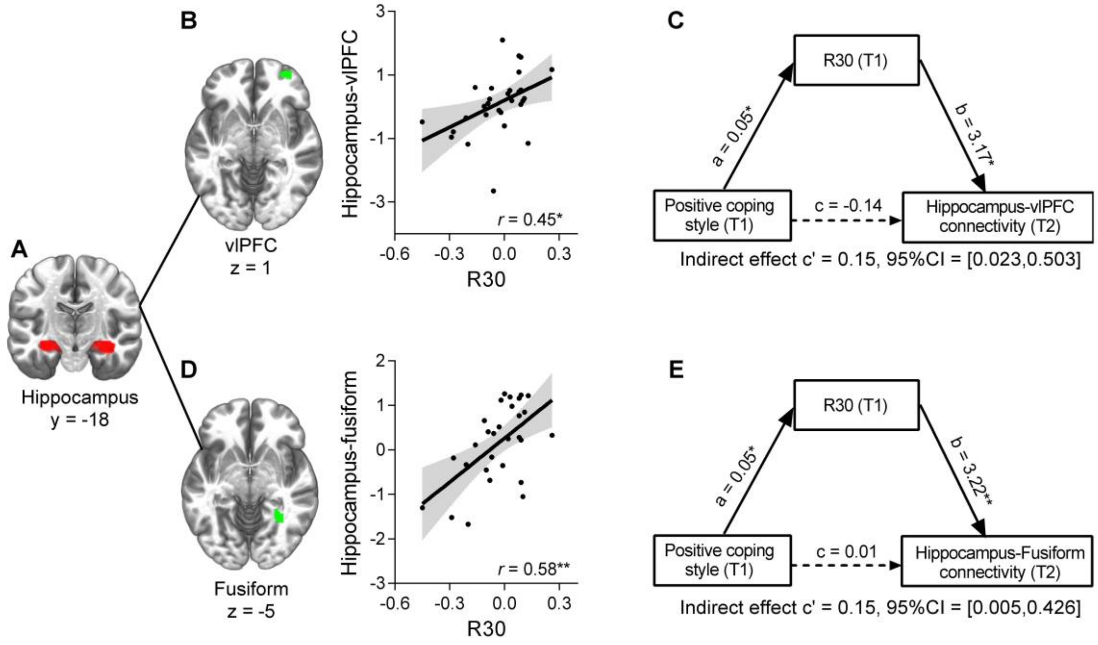


**Figure S6. The mediation models of PCS-R30-hippocampal connectivity in the second year.** (A) A representative coronal slice of the seed region of bilateral hippocampus. (B) A positively significant relationship between the first-year R30 and the second-year hippocampus-fusiform connectivity, as well as (D) the relationship between R30 (T1) and the hippocampus-vlPFC connectivity. (C) A mediation model indicating the mediation effect of the first-year R30 on the relation between the first-year PCS and the hippocampus-vlPFC (T2) connectivity as well as (E) the hippocampus-fusiform functional connectivity (T2). R30, response within 30 minutes; T1, time 1; T2, time 2; vlPFC, ventrolateral prefrontal cortex; *, $p < 0.05$; **, $p < 0.01$; 95% CI, 95% confidence interval.

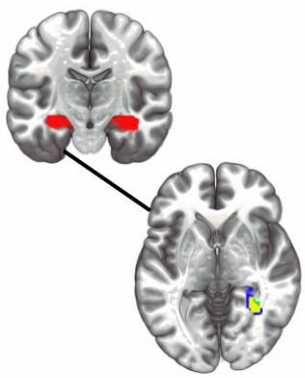


**Figure S7. A schematic view of locations of hippocampus seed-based connectivity with the fusiform areas.** The upper panel: the seed region of bilateral hippocampus (upper red brain area). The lower panel: the lower brain regions show three locations of fusiform: the green fusiform region showing the second-year fusiform region significantly related to the first-year R30, the blue fusiform region showing the change of the hippocampus-fusiform functional connectivity over one year was significantly related to the first-year R30, while the yellow area is the common fusiform region between the green fusiform region and the blue fusiform region.

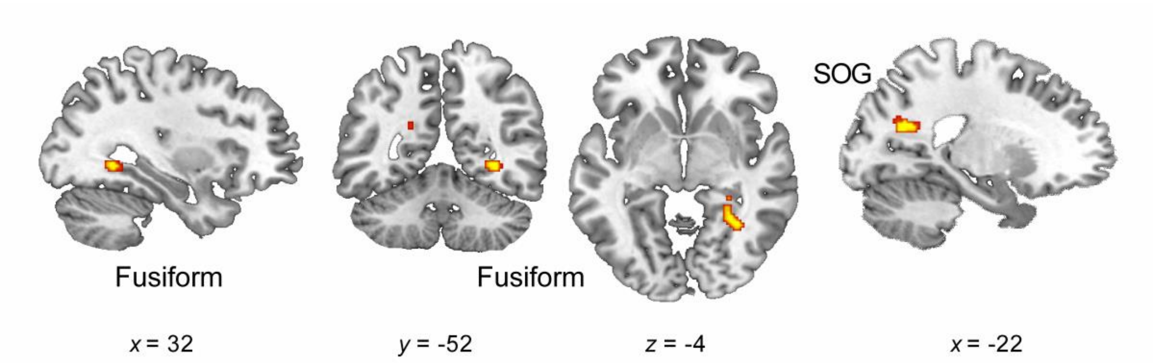


**Figure S8. Target brain regions with significant changes in functional connectivity with the hippocampus seed (T2-T1) that were related to children's R30.** SOG, superior occipital gyrus.

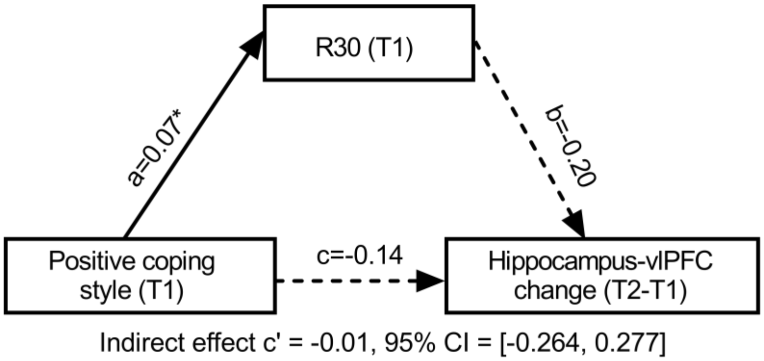


**Figure S9. R30 not mediating in the relation between the PCS (T1) and the development of hippocampus-vlPFC connectivity (T2-T1).** T1, Time 1, the first year; T2, Time 2, the second year; vlPFC, ventrolateral prefrontal cortex; 95% CI, 95% confidence interval.
